# Asymmetric stem cell division maintains genetic heterogeneity of tissue cells

**DOI:** 10.1101/2024.05.16.594576

**Authors:** Muhammed Burak Bener, Boris M. Slepchenko, Mayu Inaba

## Abstract

Within a given tissue, the stem cell niche provides the microenvironment for stem cells suitable for their self-renewal. Conceptually, the niche space constrains the size of a stem-cell pool, as the cells sharing the niche compete for its space. It has been suggested that either neutral-or non-neutral-competition of stem cells changes the clone dynamics of stem cells. Theoretically, if the rate of asymmetric division is high, the stem cell competition is limited, thus suppressing clonal expansion. However, the effects of asymmetric division on clone dynamics have never been experimentally tested. Here, using the *Drosophila* germline stem cell (GSC) system, as a simple model of the in-vivo niche, we examine the effect of division modes (asymmetric or symmetric) on clonal dynamics by combining experimental approaches with mathematical modeling. Our results suggest that the rate of asymmetric division positively correlates with the time a stem cell clone takes to expand. Taken together, our data suggests that asymmetric division is essential for maintaining the genetic variation of stem cells and thus serves as a critical mechanism for safeguarding fertility over the animal age or preventing multiple disorders caused by the clonal expansion of stem cells.

## Introduction

In a tissue, stem cells reside in the specialized microenvironment for stem-cell maintenance called a niche. Within the niche, stem cells can divide both symmetrically and asymmetrically, and the proportion of these two division modes greatly varies among different stem cell systems[1]. Typically, a defect in asymmetric cell division (ACD) can lead to loss or gain of the number of stem cell and subsequent replacement or removal events mediated by symmetric cell division (SCD). So long as the replaced cells are genetically or phenotypically equivalent to the original stem cells, tissue defects may not occur. However, once a stem-cell clone gains a harmful mutation, the expansion of such a clone may lead to disruption of tissue homeostasis and initiation of a disease. In the case of germline stem cells, it directly affects genetic diversification or fertility, thus has a huge impact on animal evolution. So far, many studies have investigated the dynamics of stem cell clones, both with computational and experimental approaches[2-7]. However, the role of ACD has not been clearly demonstrated.

It has been shown that defects in ACD can cause dysregulation of cell fate determination leading to tumorigenesis [8-14]. On the other hand, ACD has also been shown to have tumor-promoting effects. For example, ACD can produce phenotypically heterogenous cell populations in cancer, which leads to resistance to treatment [15, 16]. Furthermore, the quality surveillance mechanism, such as the elimination of damaged stem cells from the tissue, relies on cell-cell competition promoted by SCD and restricted by ACD [17-19]. Thus, it is hard to say in advance that the impact of ACD on tissue homeostasis will be definitively positive or negative, as it may vary significantly, depending on ACD/SCD rates, proliferation rates, and sizes of individual stem cell pools, that need to be independently evaluated.

Studying the *Drosophila* GSC system has yielded valuable information for our understanding of many conserved stem-cell characteristics. At the tip of the *Drosophila* testis, 8-10 GSCs adhere to a cluster of somatic cells called the hub, a major niche component. The hub cells send signals which are vital for GSC maintenance. In this microenvironment, GSCs typically divide asymmetrically via a stereotypical spindle orientation. This ACD results in only one daughter cell maintaining attachment to the hub and retaining its stem cell identity, and a second daughter cell is displaced away from the hub and becomes a gonialblast (GB) that begins to differentiate[12, 20] (Figure 1A).

**Figure 1.**
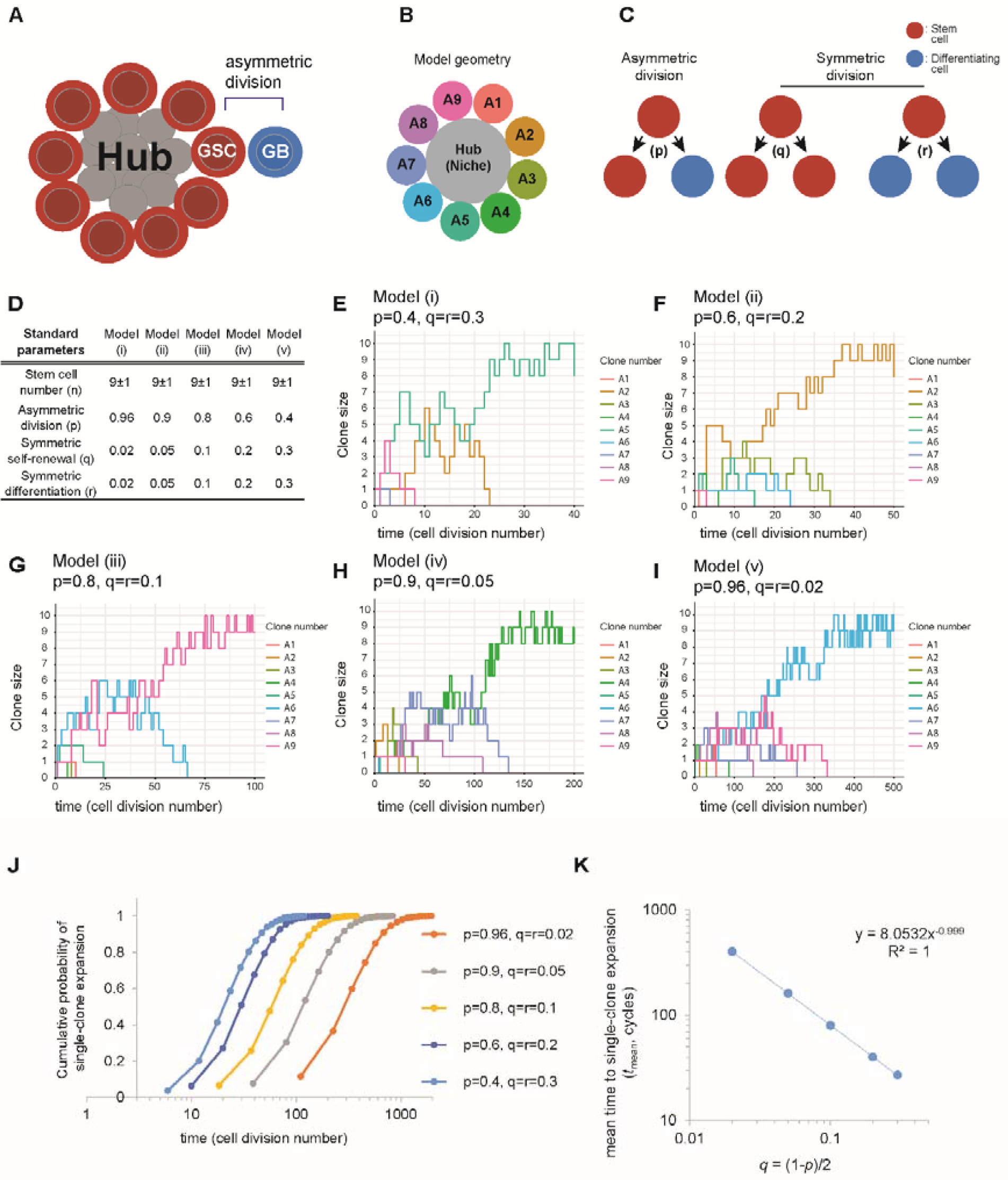
The effect of asymmetric cell division on stem cell clonality. **A)** Schematic of the hub in the *Drosophila* testis. **B)** Hub geometry reflected by the model. Stem cell clones are labeled A1 through A9. **C)** Illustration of three potential outcomes of a stem cell division: asymmetric cell division (*p*), symmetric self-renewal (*q*) and symmetric differentiation (*r*). **D)** Standard parameters for models i through v, including stem cell number (*n*) and division probabilities (asymmetric: *p*, symmetric: *q* and *r*). **E-I)** For each model (i through v), a single representative system trajectory that most closely matched the median number of cell cycles required for a single clone to dominate the niche. **J)** Cumulative probability graph of single-clone expansion over the number of cell cycles. **K)** The model yields an inverse relationship between the mean time to single-clone expansion and the rate of symmetric self-renewal ().

Owing to its simple and well-characterized tissue anatomy and defined cellular markers, this model is especially suited to addressing the impact of ACD defects on the clonal expansion rate of stem cells using both computational and experimental approaches.

## Results

### Asymmetric cell division delays the clone expansion

To assess the role of ACD in the clonal expansion of stem cells in normal tissue homeostasis, we simulated dynamics of stem-cell clones in the testicular niche using *Virtual Cell* (VCell), a publicly available software for computational modeling biological processes [21, 22]. Our model mimics real GSC niches in *Drosophila* testes. Given that the GSC number is strictly controlled depending on available niche space, the capacity of the niche in the model is constantly adjusted to maintain 9 (plus/minus one) cells that are dividing asynchronously every 12 hours (Figure 1B), (see section *Mathematical modeling* for details).

In what follows, the niche occupied by a single clone is termed “a clonal niche”. We simulated time-dependent probabilities of emergence of a clonal niche for varying frequencies of ACD (*p*) and SCD (symmetric self-renewal (*q*), and symmetric differentiation (*r*)). These frequencies satisfy the following constraints: *p* + *q* + *r* = 1 per cycle, and *q* = *r* (Figure 1C). Each stem cell clone, labelled A1 through A9, was assigned identical values of *p, q*, and *r*, and represented by unique colors in the simulation outputs. We ran 3,000 simulations for each of the parameters sets described in Figure 1D and recorded the number of each stem cell clone after every cell division event occurring in the virtual niche, see results of representative single trials in Figure 1E-I. The cumulative probabilities of single-clone expansion shown in Figure 1J demonstrate how the frequency of ACD (*p*) influences the clonal dynamics within the niche.

Strikingly, our simulation results revealed the positive correlation between the time required for emergence of a clonal niche and the rate of ACD (Figure 1K). The model demonstrates that ACD has a delaying effect on the onset of clonal expansion of stem cells.

### Fitting the model to experimental results using *Drosophila* germline stem cells

We conducted short-term and long-term lineage tracing experiments in *Drosophila* testis to monitor the clone dynamics *in vivo*. To permanently mark the GSC clones by GFP, we induced Flippase expression (hs-FLP) by incubating flies in 37 degrees for short time (heat shock), which removes the stop codon located between nanos promoter and Gal4, thereby initiating the UAS-GFP expression (Figure 2A) [23]. We optimized the heat-shock duration to 40 minutes to mark approximately 50% of the GSCs (Figure 2B). We then followed the frequency of GFP-positive GSCs at days 7, 14, and 21 post-marking. If GSCs divided only asymmetrically (*p* = 1), the GFP-positive clone frequency would remain around 50%. However, in the presence of symmetric events, the clonal expansion will cause the GFP-positive GSCs either to expand or vanish, so their frequency will tend to 0% or 100% (Figure 2C). During our imaging period, we indeed observed a shift of the GFP-positive GSC frequency towards all or none (Figure 2D). As we showed in the previous section, the time to clonal expansion is linked to *p*. Thus, how the distribution of GFP-positive GSC frequencies changes with time should tell us about the ACD rate.

**Figure 2.**
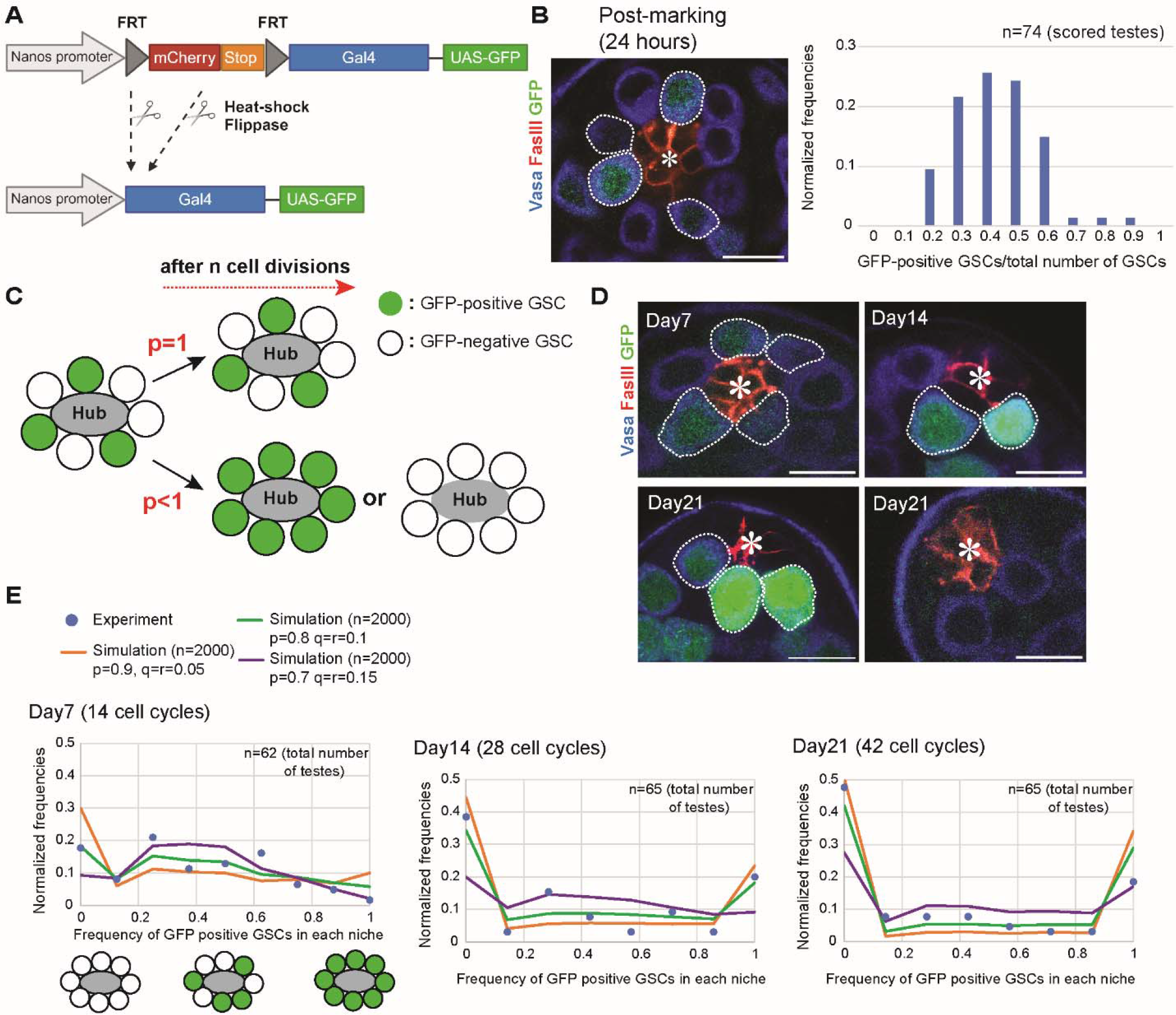
Long-term clonal tracing of germline stem cells. **A)** Illustration of the marking system. The heat-shock driven UAS-FLP expression removes the mCherry-stop cassette, and activate the expression of GFP in GSCs under the control of nosGal4. **B)** Representative immunofluorescence image of the testicular niche 24 hours after a 40-minute of heat-shock. The graph shows the marking efficiency. Note that, the testes with no marked GSCs were excluded from the day0, and a proportionate number of unmarked testes were similarly excluded from the subsequent time points (day7, 14 and 21) (see “Mathematical modeling” in the Method section for details). **C)** Possible outcomes of population dynamics of GFP positive stem cells after marking 50% of GSCs. If GSCs exclusively undergo asymmetric division (*p*=1), the frequency of GFP-positive clones would **C)** remain around 50%. Conversely, if symmetric events occur, the distribution of GFP-positive GSCs within each testis would tend towards 0% or 100%. **D)** Representative immunofluorescence images of the testicular niche at day7, 14 and 21. The upper panel shows images where the proportion of GFP-positive GSCs reaches 100%. In contrast, the lower panel shows images where the GFP-positive GSCs decline to 0%. **E)** Fitting the model to the experimental data. Simulation results are from 2,000 runs for each set of parameters: 1) *p* = 0.9, *q = r* = 0.05, 2) *p* = 0.8, *q= r* = 0.1, and 3) *p* = 0.7, *q = r* = 0.15. Root-mean-square deviations (RMSD) of the fits from the experimental data are indicated on the graph. The “n” indicates the number of imaged testes. All scale bars correspond to 10 μm. Asterisks in the images indicate the approximate location of the hub (niche).

To accurately determine the ACD rate *p* (as well as values and *r*), we have fitted our model to the experimental data, see section *Utilizing the mathematical model for analyzing experimental data* for details. We ran 2,000 simulations for each of the following parameter sets: 1) *p* = 0.9, *q = r* = 0.05; 2) *p* = 0.8, *q = r* = 0.1, and 3) *p* = 0.7, *q = r* = 0.15, and found that the best fit to our experimental results was provided by *p* = 0.8, *q = r* = 0.1 (all per cycle) (Figure 2E). This result is consistent with our prior data [24] and a previous study [25].

Our finding, however, differs from the estimate *p* ∼ 0.94-0.98 per cycle reported previously by Salzmann et al. [23]. They used short heat shocks to generate a small number of GFP-positive cells per niche and classified them as singlets, doublets, etc., depending on their mutual positions in the niche (Figure 3A). The authors obtained their estimate by interpreting multiplets as outputs of symmetric self-renewal and analyzing how the fractions of singlets were changing over several cell cycles.

**Figure 3.**
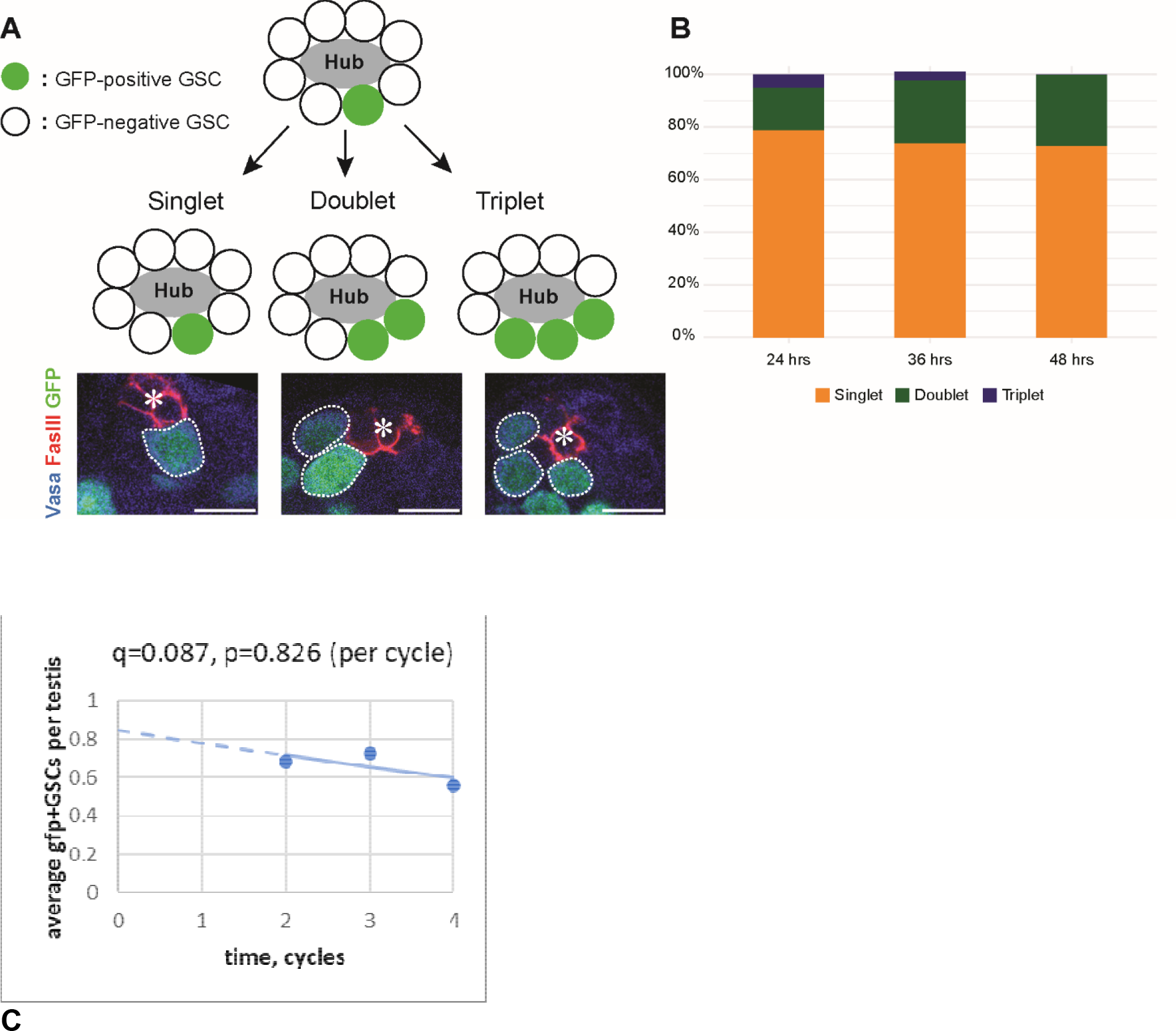
Short-term lineage tracing of germline stem cells. **A)** Examples of singlet, doublet, and triplet GSCs 24 hours post-heat-shock treatment (30 mins) that has been optimized to mark ∼1 GSC per testis. Representative immunofluorescence images of the singlet, doublet, and triplet examples. The singlet is observed when the marked GSC divides asymmetrically. The doublet and triplet are observed when the marked GSC undergoes symmetric self-renewal. **B)** The graph illustrates the percentage of singlet, doublet, and triplet GSCs at 24-, 36- and 48-hours post-heat-shock treatment. **C)** Average number of all GFP-positive GSCs per niche as a function of time. The trendline is a single exponential *Ae*^−*kt*^, where the rate constant *k* ≈ 0.087 per cycle is a rough estimate of *r*, yielding *p = 1 − 2r* ≈ 0.826 per cycle (*A* ≈ 0.85). The “n” indicates the number of imaged testes. All scale bars correspond to 10 μm. Asterisks in the images indicate the approximate location of the hub (niche).

This approach, however, is prone to underestimating the net decrease in the number of singlets. Shortly after the shock, adjacent singlets might be misinterpreted as doublets, whereas at later times, an output of self-renewal, which in *Drosophila* testes occurs mostly by way of dedifferentiation [24], might be mistaken for two individual singlets, since the reattachment of a dedifferentiated cell to the hub can occur at any position; together, these errors would cause the underestimation of the net drop in the number of singlets. We confirmed this by performing similar experiments (Figure 3). The singlet effects of identification (Figure 3B) yielded a decrease in the fraction of singlets of ∼6% over 2 cell cycles, resulting in *p* ≈ 0.94 per cell cycle, close to the estimates by Salzmann et al., but when we looked at the effects of symmetric divisions on the number of all GFP-positive GSCs, thus avoiding the potential pitfalls of discriminating singlets from multiplets, a rough estimate based on a single-exponential fit yielded *p* ≈ 0.83 per cell cycle (Figure 3C), much closer to the estimate, which we obtained by exploiting the dependence of the clonal expansion on the frequency of ACD.

## Discussion

Tissue stem cells reside in specialized microenvironments called the niche. Traditionally, it has been believed that asymmetric division is the primary mechanism for stem cell maintenance by balancing self-renewal and differentiation. However, the niche is constrained in size, leading to continuous competition among stem cells for space through a neutral drift. Consequently, the stem cells can also be maintained through symmetric divisions without expansion or reduction. In most tissues, both asymmetric and symmetric divisions contribute to stem cell maintenance, but the role of asymmetric divisions in the long-term tissue homeostasis has often been overlooked.

In this study, we found that a niche with a high probability of ACD (*p* > 0.9) never reaches a clonal results niche (the condition in which a niche is occupied by a single clone) within the average lifetime of flies (∼120 divisions, equivalent to ∼60 days) [26, 27]. In contrast, a low probability of ACD (*p* < 0.4) in a niche that quickly becomes clonal (∼22 divisions, 11 days, Figure 1E-I), supporting our hypothesis that ACD is an essential factor for preventing clone expansion.

Our findings provide significant insights into clonal dynamics of stem cells and suggests potential therapeutic strategies that could manipulate the division modes of stem cells to suppress clonal expansion in disease conditions. The mathematical model of clonal expansion, used in this study to obtain accurate estimates of the rates of ACD and SCD in *Drosophila* testes, could also be adapted to predict stem cell behavior in mammalian stem cell niches, across both steady-state and various pathological conditions.

In recent years, clonal expansion of stem cells has emerged as a crucial factor in the development of multiple pathological conditions. Clonal hematopoiesis (CH) is a condition characterized by the presence of clonally expanded hematopoietic stem cells (HSCs) and has emerged as a risk factor for hematopoietic malignancy and cardiovascular diseases [28, 29]. Extensive clinical studies have identified multiple commonly mutated genes within these expanding HSC clones and explored the mechanisms by which these mutations contribute to disease progression [28, 30]. Removal of mutant clones by manipulating competition between mutant and normal cells has been suggested as a novel strategy of definitive treatment [31-38], but much of mechanistic knowledge required for implementation of such treatment is still lacking.

## Materials and methods

### Fly husbandry and strains

All fly crosses were performed on standard Bloomington medium at 22°C to avoid hs-flp expression and indicated age (days after clone marking) of flies were used for all experiments. The following fly stocks were obtained from Bloomington stock center (BDSC); *hs-flp* (BDSC1929). nos-FRT-mCherry-stop-FRT-Gal4, UAS-GFP [23] was from Inaba lab stock.

### GFP labeling of germline stem cells

Flies carrying the heat-shock-FLP allele were crossed with nos-FRT-mCherry-stop-FRT-Gal4, UAS-GFP flies and maintained at 22°C. 0-3 days old progenies were heat-shocked in a 37°C water bath once for 40 min (optimized for 50% marking) in vials with fly food. Vials were placed back at 22°C and testes were dissected at desired time points (24hrs, day7, 14 and 21 post-heat shock).

### Immunofluorescence Staining

Testes were dissected in phosphate-buffered saline (PBS) and fixed in 4% formaldehyde in PBS for 30–60 minutes. Next, testes were washed in PBST (PBS + 0.2% TritonX-100, Thermo Fisher) for at least 60 minutes, followed by incubation with primary antibody in 3% bovine serum albumin (BSA) in PBST at 4°C overnight. Samples were washed for 60 minutes (three times for 20 minutes each) in PBST, incubated with secondary antibody in 3% BSA in PBST at room temperature for 2 hours and then washed for 60 minutes (three times for 20 minutes each) in PBST. Samples were then mounted using VECTASHIELD with 4’,6-diamidino-2-phenylindole (DAPI) (Vector Lab).

The primary antibodies used were as follows: rat anti-Vasa (1:20; developed by A. Spradling and D. Williams, obtained from Developmental Studies Hybridoma Bank (DSHB); mouse anti-Hts (1B1, 1:20; DSHB) mouse-anti-FasIII (1:20, 7G10; DSHB); AlexaFluor-conjugated secondary antibodies (Abcam) were used at a dilution of 1:400.

### Mathematical modeling

To study dynamics of unsynchronized clones, we formulated a continuous-time stochastic model. We first assume that all clones are described by same probability rates of outcomes of randomly occurring asymmetric and symmetric divisions, thus, there are three model parameters: the rate constant of asymmetric division *p*, and the rate constants of symmetric self-renewal *q* and symmetric differentiation *r*.

Let *v* be the division frequency, then *p* + *q* + *r* = *v*. Because cell cycle duration of stem cells sets the model time scale, it is convenient to use it as a time unit, then 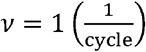 and 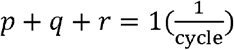. Since the average number of cells in the niche is constant, then *q* must equal *r*, so the system has only one independent parameter. Note that even with *q = r*, fluctuations of the niche population around its average size would grow with time, unless additional regulation is exercised, as described below.

Our model is formulated in terms of numbers of descendants of individual clones {*n*_*i*_}, where index *i* enumerates original clones (and their descendants): *i* = 1, …, *N*, where *N* is the niche capacity. The dynamics of *n*_*i*_ are governed by the randomly occurring events of asymmetric and symmetric divisions.

To restrict fluctuations of the population size, the niche must ensure that the total number of cells, 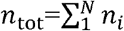, at any time remains close to *N* : |*n*_tot_ − *N* < ε where is a small positive integer. Care must be taken to interpret such control in a biologically consistent manner (see detailed discussion later in this section). Our implementation of the tight control over *n*_tot_ by the niche assumes strong correlation between the proliferation and differentiation occurrences. Specifically, the system is governed by the following transition probabilities:

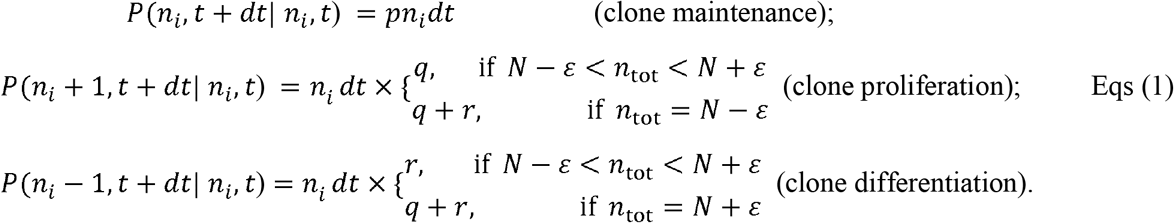

The model was solved by generating stochastic trajectories of the system in VCell [21] with a nonspatial Monte Carlo solver based on the Gibson - Bruck method [39]. The clonal expansion was analyzed for the uniform initial conditions, *n*_*i*_ | _*t*=0_ = 1for all *i* = 1, …, *N*. Results indicate that our model retains features of discrete-time Moran-type stochastic processes [40, 41], with all *n*_*i*_ being interdependent.

Note that the inequality | *n*_tot_ − *N* | ε can also be enforced by lowering transition probabilities of the symmetric outcomes that violate this inequality or setting such probabilities to zero. Below is an example of postulating low probabilities of unwanted symmetric outcomes:

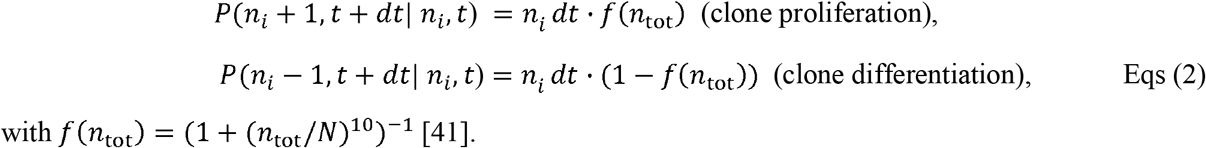

with *f*(*n*_tot_) = 1 + (*n*_tot_/*N*)^10^)^−1^ [41].

Alternatively, the probabilities of the outcomes violating the limits for *n*_tot_ could be set to zero:

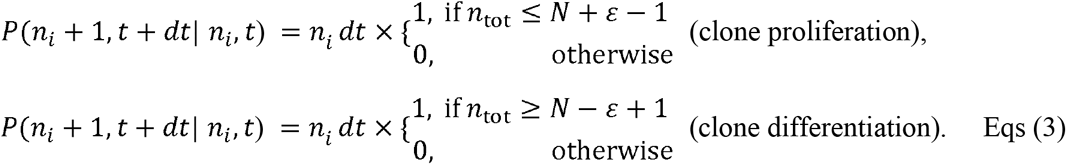

(The transition probability of asymmetric divisions in these approaches is the same as in Eqs (1)).

These models, however, result in an artificial dependence of a cell cycle period on the type of division the stem cell decides to undergo. Indeed, because these methods effectively rule out certain symmetric outcomes, there are on average fewer symmetric divisions per unit time than asymmetric ones. As a result, the average period of a symmetric division is longer than that of an asymmetric one.

Such an artefact does not arise in the model defined by Eqs (1).

### Utilizing the mathematical model for analyzing experimental data

We used the mathematical model described in the previous section for analyzing data obtained in experiments with transgenic male flies that enable induction of GFP-labeled stem cell clones. The experiments were aimed at determining values of *p, q* and *r* characteristic of the population dynamics of GSCs in *Drosophila* testes.

Frequencies of symmetric divisions in *Drosophila* testes were estimated in an earlier study [23], where observed population dynamics of GFP-marked cells were also compared against predictions from a mathematical model. The model used in [23] was applicable only to experiments with few GFP-positive cells per niche at any time, because it ignored dynamics of unmarked cells^1^. However, amassing statistically significant data from experiments with small number of GFP-positive cells per niche requires examining large pools of flies, which makes these experiments inefficient.

In our experiments, the flies were inducted to have initial numbers of marked stem cells of about half the niche capacity. As in [5], we tracked over time the fractions of GFP-positive cells in the niches, 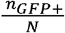. Specifically, four portions of a large pool of inducted flies were sacrificed at times separated by seven days, with time zero being the closest to the time of induction. For each time point, testes were dissected, the total number of stem cells and the number GFP-positive cells were counted, and their ratio 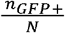 was computed for each testis. The value of *p* was determined by comparing experimental distributions of 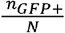 at the specific times *t*, 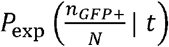, to corresponding theoretical distributions 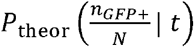 predicted for varying *p* our model whose applicability was not limited to small *n*_*GFP* +_.

We derived the predicted distribution from multiple stochastic trajectories generated by a Monte Carlo simulator according to Eqs (1) (see the previous section). In a single run, the solver returns an instance of *n*_*GFP* +_ at time *t* for a given *N* and an initial value of *n*_*GFP* +_ (in all calculations, we used ε = 1 and assumed that *N* does not change with time). Because in the experiments *N* varied from subject to subject, the predicted distribution 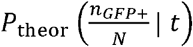 is a weighted average of the distributions obtained for each *N*, experimentally observed at time *t*, and each *n*_*GFP* +_ observed at *t* = 0, which did not exceed the given *N*.

Accordingly, we first determined, by statistically processing the experimental results for each time point *t*, the frequencies, or ‘weights’, of all values of *N* observed at time *t,w*_*exp*_ (*N,t*). Similarly, we determined the frequencies of all initial *n*_*GFP* +_ for each *N* observed at time *t, w*_*exp*_ (*n*_*GFP* +_,0 | *N,t*). Both weights were normalized to unity.

Secondly, for each pair of *N* and initial *n*_*GFP* +_, we determined the probability distribution of 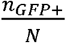 at a time point *t*, 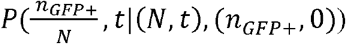. For this, we simulated large amounts of predicted *n*_*GFP* +_ by running the Monte Carlo solver with Eqs (1). Then the predicted distribution was computed as

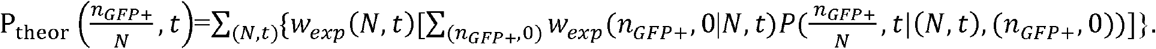

Note that one stochastic trajectory of the model with the uniform initial values: *n*_*i*_ | _*t*=0_ =1 for all *i* = 1, …,*N*, yields 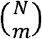 instances of *n*_*GFP* +_ at time *t*, where *m* denotes the number of the GFP-positive cells at *t* = 0, 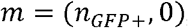, and 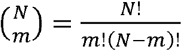. This is because every combination of *m* variables out of the total of *N* yields one instance of *n*_*GFP* +_ at time *t*. Alternatively, the model could be equivalently rewritten in terms of only two variables {*n*_0_, *n*_1_}denoting, respectively, the numbers of the GFP-negative and GFP-positive cells in the niche. Eqs (1) still apply, and *N* retains the meaning of niche capacity, but *i* = 1,2 and *n*_tot_ = *n*_0_ + *n*_1_. For this version of the model, one stochastic trajectory yields exactly one instance of *n*_*GFP* +_ at time *t*.

For each time point and for each tested value of parameter *p*, we compared 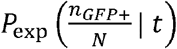 and 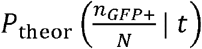 by computing the root mean squared deviations,

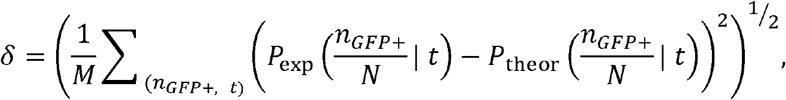

where *M* is the number of values of 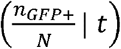 for which the probabilities were computed.

### Statistical analysis and graphing

No statistical methods were used to predetermine sample size. The experiment values were not randomized. The investigators were not blinded to allocation during experiments and outcome assessment. Statistical analysis and graphing were performed using GraphPad prism9. All data are shown as means ± s.d. The adjusted P values from Šidák’s multiple comparisons test are provided; shown as *P<0.05, **P<0.01, ***P<0.001, ****P<0.0001; NS, non-significant (P≥0.05).

## Supplemental Data

Individual numerical values displayed in all graphs are provided.

## Acknowledgements

We thank Yukiko M. Yamashita and the Bloomington *Drosophila* Stock Center and the Developmental Studies Hybridoma Bank for reagents. This research is supported by R35GM128678 from National Institute for General Medical Sciences, start-up funds from UConn Health (to M.I.), and UConn Research Excellence Program (REP). BMS was supported in part by R24GM137787 from National Institute for General Medical Sciences.

## Author Contributions

All authors contributed equally for conception and design, interpretation of data and wrote and edited the manuscript. B.M.S and M.B.B performed mathematical modeling and data fitting using VCell. M.B.B and M.I. performed acquisition and analysis of experimental data.

## Declaration of Interests

The authors declare no competing interests.

## Footnotes

1 Generally, the dynamics of marked and unmarked cells are interdependent because of the niche control over the total number of GSCs, marked and unmarked, that is kept constant.

## Notes

### Competing Interest Statement

The authors have declared no competing interest.

### Summary of Updates

In this version, we revised our interpretation for difference in the frequency of asymmetric division between our study and previously published study.

